# Response to nitrogen and salinity in *Rhizophora mangle* propagules varies by maternal family and population of origin

**DOI:** 10.1101/2021.08.04.454989

**Authors:** Christina L. Richards, Kristen L. Langanke, Jeannie Mounger, Gordon A. Fox, David B. Lewis

## Abstract

Many coastal foundation plant species thrive across a range of environmental conditions, often displaying dramatic phenotypic variation in response to environmental variation. We characterized the response of propagules from six populations of the foundation species *Rhizophora mangle* L. to full factorial combinations of two levels of salt (15 ppt and 45 ppt) reflecting the range of salinity measured in the field populations, and two levels of nitrogen (N; no addition and amended at approximately 3 mg N per pot each week) equivalent to comparing ambient N to a rate of addition of 75 kg per hectare per year. The response to increasing salt included significant plasticity in succulence. Propagules also showed plasticity in maximum photosynthetic rate in response to N amendment, but the responses depended on the level of salt and varied by population of origin. Generally, survival was lower in high salt and high N, but the impact varied among populations. Overall, this study revealed significant phenotypic plasticity in response to salt and N level. Propagules from different populations differed in all traits measured. Variation in phenotypic plasticity and propagule survival in *R. mangle* may contribute to adaptation to a complex mosaic of environmental conditions and response to climate change.

## INTRODUCTION

Many plant species thrive across an extensive range of environmental conditions, often displaying dramatic phenotypic variation (McKee, 1995; Smith and Snedaker, 1995; Richards et al., 2005; Feller et al., 2010). This is particularly true in coastal ecosystems that are marked by temporal cycles and spatial variation in tidal inundation, temperature, nutrient availability, and salinity (Pennings and Bertness, 2001; Krauss et al., 2008; IPCC, 2014; Proffitt and Travis, 2014; Wuebbles et al., 2014). In addition to the naturally dynamic habitat of coastal systems, anthropogenic activities can increase the input of nutrients and alter watersheds, further contributing to environmental variation (Bertness et al., 2002; Barbier et al., 2008; Gedan et al., 2009, 2011; Crotty et al., 2020). Within these dynamic conditions, foundation plant species, like mangroves, provide ecosystem services, such as habitat for many juvenile fish species, biotic filters of pollutants, and storm buffers (Ellison et al., 2005; Zedler and Kercher, 2005; IUCN, 2007; Alongi, 2008, 2013; Costanza et al., 2008; Gedan et al., 2011). Foundation species are defined not only as those that dominate a community assemblage numerically or in biomass, but they also determine diversity of associated taxa through a variety of interactions (Ellison, 2019). Further, foundation species modulate fluxes of nutrients and energy in their ecosystem (Ellison, 2019). Hence, these species disproportionately contribute to maintaining habitat integrity and ecosystem resilience (Bertness and Callaway, 1994; Keith et al., 2017; Ellison, 2019; Bertness, 2020; Qiao et al., 2021). Understanding how these species cope with challenges from anthropogenic impacts is key to preserving the ecosystems they create and define (Guo et al., 2021).

Understanding the mechanisms of response in coastal foundation species has become increasingly important for conservation and management strategies as these species must cope with rising sea levels and increased warming due to global change (Gedan et al., 2011; Kirwan and Megonigal, 2013; Osland et al., 2013, 2017). The Food and Agriculture Organization of the United Nations (FAO) estimates that as much as 35% of global mangrove forest habitat has been destroyed since roughly 1980 for the development of human settlements, agriculture and aquaculture, and industrial shipping harbors, although the rate of loss appears to have slowed in the last decade (Food and Agriculture Organization of the United Nations, 2007; Polidoro et al., 2010; Ellison et al., 2015; Global forest resources assessment 2020, 2020). In some regions, mangrove trees are also harvested for wood and charcoal (Ellison et al., 2015), resulting in habitat fragmentation and isolation of existing remnant fragments (Friess et al., 2012; Haddad et al., 2015). The resultant loss of diversity could pose risks for these coastal foundation species in the future, particularly as sea levels are projected to rise between 0.2 and 2 m over the next century due to anthropogenic climate change (Melillo et al., 2014).

The vulnerability of coastal foundation plant communities to global change has been debated. Several authors have suggested that the combination of eutrophication and sea-level rise may result in synergistic losses and requires further research (Deegan et al., 2012; Kirwan and Megonigal, 2013; Kirwan et al., 2016; Crosby et al., 2017; Schuerch et al., 2018). While coastal eutrophication may enhance growth of foundation species, nutrient enrichment studies report a range of impacts on coastal systems depending on the local conditions (Anisfeld and Hill, 2012; Kirwan and Megonigal, 2013). Local sediment characteristics, soil nutrients, microbial processes, and shifts in allocation of the plant species can impact future nutrient cycling and marsh stability (McKee et al., 2007; Turner, 2011; Deegan et al., 2012; Lewis et al., 2021).

Predicting species level responses to environmental challenges requires an understanding of the amount of phenotypic variation within and among populations, which may reflect both phenotypic plasticity and heritable differences in phenotype (Richards et al., 2006; Nicotra et al., 2010; Banta and Richards, 2018). Several studies have shown that species harbor heritable differences in eco-physiological traits (Arntz and Delph, 2001; Geber and Griffen, 2003; Caruso et al., 2005), and variation in plasticity of traits (Sultan, 2001; Matesanz and Sultan, 2013; Nicotra et al., 2015; Matesanz et al., 2021). However, the amount of variation in natural populations for traits that are important for response to future climates is not well known (Davis and Shaw, 2001; Parmesan, 2006). Plants that inhabit coastal and intertidal zones have putative physiological adaptations that enable them to grow and reproduce in the anoxic and saline conditions that characterize these habitats. These adaptations include reduction of water required by the plant, adjustment of carbon uptake and nutrient absorption, and changes in resource allocation (Antlfinger and Dunn, 1979; Cavalieri and Huang, 1979; Antlfinger and Dunn, 1983; Glenn and O’Leary, 1984; Donovan et al., 1996, 1997; Flowers and Colmer, 2008). In addition, mangrove species can moderate anoxia that results from flooding via root growth, and altered peat formation has allowed mangrove communities to keep pace with sea level rise (McKee et al., 2007). Mangroves have been shown to respond to changes in nitrogen (N) by altering relative growth rate, photosynthetic rate, and resource allocation (Feller, 1995; McKee, 1995; Feller et al., 2003), which could be an important response to anthropogenic activities, such as runoff from agriculture and other types of land use change (Feller et al., 2003; Alongi, 2013).

Given spatial differences in salinity, anoxia, and N in the intertidal habitat, plasticity of traits that allow for tolerating such conditions may be adaptive. We expect intertidal plants like mangroves to show plasticity in response to salinity and N conditions. Profitt & Travis (Proffitt and Travis, 2010) found plasticity in growth rate and reproductive output within and among natural *Rhizophora mangle* mangrove populations in the Tampa Bay region. However, they also found both site of origin and maternal tree of origin affected *R. mangle* growth and survival, and that these effects varied by intertidal position (significant maternal family by elevation interaction; (Proffitt and Travis, 2010)). On the other hand, we found that nearby populations had low genetic diversity, and little population differentiation. Instead we discovered high epigenetic diversity based on DNA methylation polymorphisms (Mounger et al., 2021). This type of epigenetic diversity has been associated with phenotypic and functional diversity, and could be a mechanism underlying phenotypic plasticity (Zhang et al., 2013; Nicotra et al., 2015; Herrera et al., 2017; Jueterbock et al., 2020). Several studies suggest that epigenetic diversity could be particularly important in genetically depauperate species, providing a nongenetic source of response to the diverse conditions experienced by these populations (Gao et al., 2010; Verhoeven et al., 2010; Richards et al., 2012; Jueterbock et al., 2020). Further, some epigenetic differences have been shown to be heritable. In fact, we discovered that the differences in DNA methylation in *R. mangle* propagules were predicted by maternal tree suggesting a high degree of heritability of differences in DNA methylation (Mounger et al., 2021).

In this study, we characterized within and among population level variation in putative adaptive traits in response to combinations of salinity and N in a full factorial design. Given the dynamic environment inhabited by *R. mangle* and the evidence of heritable nongenetic differences among populations, we predicted differences in response to salinity and N amendment treatments among propagules collected from different populations. Our study was designed to test three predictions. First, *R. mangle* seedlings will be plastic in response to salinity and N amendment in putative adaptive traits that conserve water and adjust allocation of N. Second, response to salinity and N amendment will co-vary as plants shift resources to maintain osmotic balance. Finally, populations will vary in putative adaptive traits, and in plasticity of these traits, due to population differentiation.

## MATERIALS AND METHODS

### Study species

The red mangrove, *Rhizophora mangle* L. 1753 (Malpighiales, Rhizophoraceae), is an evergreen shrub or tree found along tropical and subtropical coastlines across the Americas, East Africa, Bermuda, and on a number of outlying islands across the South Pacific (Proffitt and Travis, 2014; Tomlinson, 2016; DeYoe et al., 2020) that can grow to heights of twenty-four meters (Bowman, 1917). Poleward expansion of the species is limited by freezing events (its current northern range limit is roughly 29° N latitude; (Proffitt and Travis, 2014; Kennedy et al., 2017). It is considered a self-compatible species, with selfing rates in Tampa Bay estimated to be as high as 80-100% (Proffitt and Travis, 2005; Nadia and Machado, 2014). However, colder temperatures and contaminants from anthropogenic sources correspond with increased flowering and outcrossing, potentially resulting in higher genetic diversity particularly in the smaller populations at range limits from 28-30° N (Proffitt and Travis, 2005, 2014). *Rhizophora mangle* stands in our study area have a mean number of about 600 reproducing trees per kilometer of estuary (Proffitt and Travis, 2014). Pollinated *R. mangle* flowers mature in approximately 95 days, producing the buoyant hypocotyl also known as a propagule (Raju Aluri, 2013). The viviparous propagule germinates and matures on the maternal tree before it drops off, is dispersed pelagically, and becomes established as a seedling (McKee, 1995). *Rhizophora mangle* excludes salt in the root system through selective uptake of potassium (K^+^) to sodium (Na^+^) ions (Wise and Juncosa, 1989; Flowers and Colmer, 2008; Krauss et al., 2008; Medina et al., 2015), and allocates resources to manage osmotic potential (Bowman, 1917; Flowers and Colmer, 2008).

### Sampling design

We sampled six populations of *R. mangle* between June 9 and June 26 of 2015, on the west coast of central Florida (USA) within the following county and state parks: Anclote Key Preserve State Park (AC), Fort De Soto Park (FD), Honeymoon Island State Park (HI), Upper Tampa Bay Conservation Park (UTB), Weedon Island Preserve (WI), and Werner-Boyce Salt Springs State Park (WB) (Figure 1). The sites varied in salinity, mean tidal range, and neighboring species. We measured salinity at each site with a refractometer finding that salinity ranged from 20 to 40 parts per thousand (ppt) across the sites at the time of collection. This area has a humid subtropical climate with mean monthly temperatures ranging from 15.6 °C in January to 28.5 °C in August (1991-2020 monthly normals, U.S. NOAA National Centers for Environmental Information, station GHCND:USC00088824, Tarpon Springs, Florida), and annual precipitation of 1379 mm (annual mean 1991-2020, Tarpon Springs station). Precipitation falls as rain, with 60 % falling during June through September, and 40 % evenly distributed among other months. Monthly mean relative humidity ranges from 67 % in April and May to 76 % in August and September (1948-2018, U.S. NOAA Comparative Climate Data for the United States through 2018, station GHCND:USW00012842, Tampa International Airport). Tides are semi-diurnal, with 0.57 m median amplitudes (U.S. NOAA National Ocean Service, Clearwater Beach, Florida, station 8726724). Sea-level rise is 4.0 ± 0.6 mm per year (1973-2020 trend, mean ± 95 % confidence interval, NOAA NOS Clearwater Beach station).

**Figure 1:**
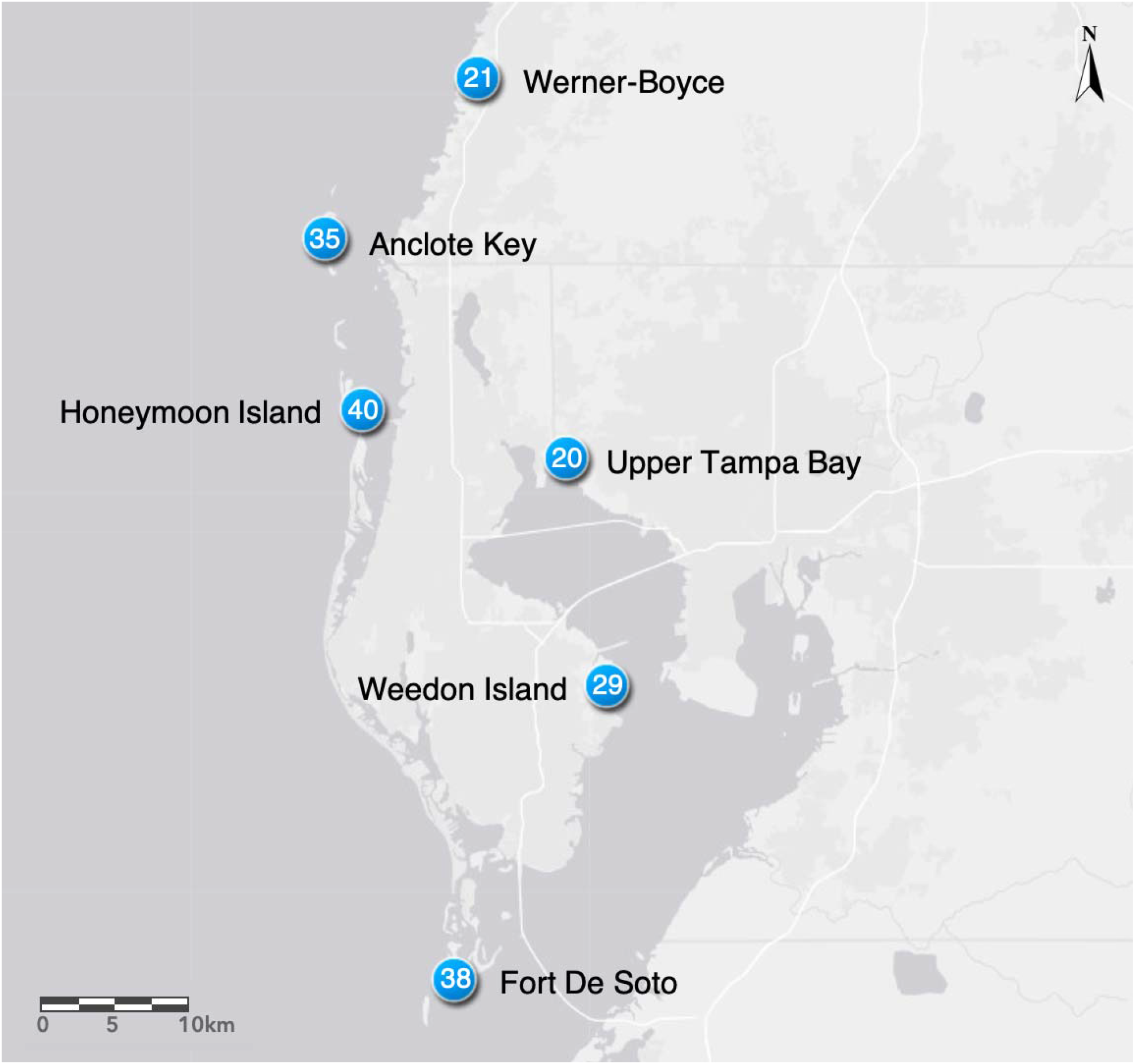
We collected *Rhizophora mangle* propagules from the following sites: Werner-Boyce Salt Springs State Park, Anclote Key Preserves State Park, Honeymoon Island State Park, Upper Tampa Bay Hillsborough County Park, and Weedon Island Preserve Pinellas County Park. Salinity levels (ppt) on the date of collection for each site are indicated within the site location markers.

Honeymoon Island had a near monoculture of *R. mangle* while the remaining sites contained mixtures of two other mangrove species that are common in Florida: *Laguncularia racemosa* L. and *Avicennia germinans* L.. We refer to plants from different sites as members of different populations based on our previous work which found differences among sites based on molecular markers (Mounger et al., 2021). At each population, we collected 20 propagules directly from each of 10 maternal trees separated by at least 10 m from each other to maximize the range of genetic variation sampled within each population (Albrecht et al., 2013). Propagules from each maternal tree were at least half-siblings but they could be more closely related due to the high selfing rate in the study area (Proffitt and Travis, 2005).

We refrigerated the propagules at 4 °C for up to 14 days until we planted them in the greenhouse at the University of South Florida Botanical Gardens. In the greenhouse, propagules from four of the maternal trees at AC and nine of the maternal trees at FD failed to establish, so we returned to sample propagules and maternal tissue from 8 new maternal trees at FD on August 12 and 29, and from the same original maternal trees at AC on October 17. Hence, while most of the propagules were in the greenhouse from the end of June until mid-October before they were exposed to treatments, propagules from these maternal families had less acclimation time before treatments began.

### Experimental treatments

We measured the length of each propagule and planted each in a 0.5-liter pot with a 50:50 mixture of sand and peat soil. We watered the plants each day with tap water until we started applying the salinity and N amendment in mid-October. We set up the experiment in five spatial blocks. Within each block we randomized the position of plants such that each block had one replicate of each family for each treatment combination (i.e., a full factorial randomized complete block design with N= 6 populations x 10 maternal families x 4 environmental treatment combinations [2 salinity x 2 N fertilization] x 5 blocks x 1 replicate/block = 1200 plants). The treatments were a factorial combination of low and high salt with either no addition or addition of N. The salt treatments were made with Morton solar salt (NaCl). The low salt treatment was 15 ppt and the high salt was 45 ppt, reflecting a slightly wider range of salinity than we measured in the field sites. The N treatments were made from two moles of N from urea (NH_4_Cl) and one mole from ammonium nitrate (NH_4_NO_3_) in tap water approximating 3mg N per pot each week. This level is similar to a rate of 75 kg N per hectare per year, based on an estimated rate of loss by soil erosion and water runoff from corn crop residue in the United States (Pimentel et al. 1989).

At the start of treatments, we recorded seedling initial height from the soil to end of any growth. To avoid osmotic shock, the salinity treatment was applied twice a week and gradually increased by five ppt each treatment. The low salt level (15 ppt) was reached in two weeks and the high salt level (45 ppt) in six weeks. We started N treatments after the first week (when salinity treatments were 10 ppt), and applied N once per week from October 15, 2015-May 1, 2016. We watered on non-treatment days with enough water to saturate the soil, but not flow through. Once per week, we watered with sufficient water to flow through the soil to prevent salt buildup. To determine if the N amendment was lost between treatments, we collected the flow-through leachate for a subset of eight plants, two of each combination of salt and N treatment. We analyzed leachate for total dissolved N via combustion and luminescent detection (Skalar Formacs TN analyzer, Breda, The Netherlands).

### Traits measured

We measured five traits related to salt tolerance and overall performance for each plant: change in height from beginning to end of treatments (cm) (hereafter, height growth), leaf mass per area or LMA (dry leaf mass in g / total leaf area cm^2^), succulence (grams of water in all leaves / total leaf area cm^2^), root to shoot ratio based on dry biomass, and total dry biomass (g) at harvest. In addition, we used a LI-COR 6400 to measure maximum photosynthetic rate (micromoles CO_2_ / m^2^ s) for a subset of the plants just prior to harvest. We determined that the appropriate photosynthetically active radiation (PAR) for saturation in these plants was 1000 micromoles PAR/ m^2^ s based on light curves generated from six data points from each of two plants (one low salt-no N and one high salt-high N). We then measured maximum photosynthetic rate on one plant with at least two healthy leaves for each surviving maternal line for each treatment (n= 29 low salt-no N, n= 31 high salt-no N, n=26 low salt-high N, and n= 32 high salt-high N, for total N=118 total plants). We defined healthy leaves as attached, a minimum of 50% green, and fully developed. We took maximum photosynthetic rate measurements at a CO_2_ rate of 400 micromoles/(m^2^·s) and a flow rate of 500 micromoles/s. We took the measurements on a healthy leaf from the second node of each plant after the leaf had been clamped in the LI-COR for one minute to ensure conditions had stabilized. We measured maximum photosynthetic rate for the 118 plants in random order over six consecutive days from April 23-28, 2016, between 8:30 and 11:30 in the morning.

We characterized each plant as alive, dormant, or dead at the end of the experimental treatments. We assigned plants that showed no growth and no desiccation to the dormant category. All live and dormant plants were harvested after six months of treatment. Height growth, succulence, maximum photosynthetic rate, and LMA was only analyzed for alive plants. We only used healthy leaves for succulence, maximum photosynthetic rate, and LMA. We included leaves that were attached, but not 50% green or fully developed in dry above ground biomass. We measured the biomass of above and below ground tissues of all harvested plants after the tissues were dried at 60°C until they maintained constant mass. Finally, we measured the total dry mass of leaves after drying in silica desiccant beads for a minimum of seven days to constant mass.

### Statistical analysis

We performed all statistical analyses in R, version 4.0.3 (R Core Team, 2020). All analyses reported here used either the General Linear Model (GLM) or Generalized Linear-Mixed-Model (GLMM) frameworks. We checked the residuals to assess normality on traits as appropriate; we did not transform height growth or photosynthesis (lmer and glm), but we used the log link function (glmer) for analysis of succulence, LMA, root to shoot ratio, and total biomass. We used the function lmer or glmer implemented within the lme4 package (Bates et al., 2015) to fit a series of models and identify the best fit model for each trait (Table 1). For each trait and for survival we began with a saturated model that included as fixed factors salt, N, and their interaction. For change in height, we included the covariate of height at the beginning of treatments in October. The saturated models also included random effects of block, population and maternal family nested within population. In several cases, models with random terms for block or population failed to converge (Table 1), most likely because there were relatively few blocks and populations. In these cases, we began by treating these as fixed effects. We used Anova (type III) in the package car (Fox and Weisberg, 2019) to evaluate the significance of fixed effects when they were included in the best model.

**Table 1.** Model selection. Abbreviations: Fam = maternal family; ht = height; LMA = specific leaf mass (dry mass/area); Photo = maximum photosynthetic rate; Pop = population; RTS = root:shoot biomass ratio; SNG = number of survivors in April. Terms in parentheses are random terms; (1|term) indicates that a random intercept is estimated for each term. Pop/Fam = maternal families nested within source populations. For some models, Block or Pop were fit as fixed effects if estimation as random effects failed (generally due to the small numbers of blocks and source populations). Δ_AIC_ is the difference between the saturated model (or closest model to it when saturated model was singular) and the AIC for the given model.

| | df | AIC | $\Delta_{AIC}$ |
| --- | --- | --- | --- |
| **Height (N=907)** |  |  |  |
| April ht ~ Oct ht + Salt * N + Block + (1\|Pop/Fam) | 12 | 4308.4 | 0 |
| April ht ~ Oct ht + Salt + N + Block + (1\|Pop/Fam) | 11 | 4306.8 | -1.6 |
| April ht ~ Oct ht + Salt + N + (1\|Pop/Fam) | 8 | 4301.8 | -6.7 |
| April ht ~ Oct ht + Salt + (1\|Pop/Fam) | 6 | 4298.2 | -10.24 |
| April ht ~ Oct ht + (1\|Pop/Fam) | 4 | 4297.4 | -10.98 |
| **LMA (N=818)** |  |  |  |
| LMA ~ Salt * N + (1\|Pop/Fam) + (1\|Block) | 8 | -8041.9 | 0 |
| LMA ~ Salt + N + (1\|Pop/Fam) + (1\|Block) | 7 | -8043.9 | -2 |
| LMA ~ Salt + (1\|Pop/Fam) | 6 | -8045.9 | -4 |
| LMA ~ (1\|Pop/Fam) + (1\|Block) | 5 | -8041.8 | 0.1 |
| **Succulence (N=818)** |  |  |  |
| Succulence ~ Salt * N + (1\|Pop/Fam) | 7 | -6534 | 0 |
| Succulence ~ Salt + N + (1\|Pop/Fam) | 6 | -6533.5 | 0.5 |
| Succulence ~ Salt + (1\|Pop/Fam) | 5 | -6534.5 | -0.5 |
| **Root:shoot biomass ratio (N=1073)** |  |  |  |
| RTS ~ Salt * N + (1\|Pop/Fam) + (1\|Block) | 8 | 1255.1 | 0 |
| **Total biomass (N=1073)** |  |  |  |
| Total biomass ~ Salt * N + Block + (1\|Pop/Fam) | 8 | 4869.7 | 0 |
| Total biomass ~ Salt * N + (1\|Pop/Fam) | 7 | 4867.8 | -0.1 |
| Total biomass ~ (1\|Pop/Fam) | 4 | 4864.5 | -5.2 |
| **Maximum photosynthetic rate (N=118)** |  |  |  |
| Photosynthesis ~ Salt * N + Pop + Block + (1\|Fam) | 14 | 592.9 | 0 |
| Photosynthesis ~ Salt * N + Site + (1\|Fam) | 10 | 587.7 | -5.1 |
| Photosynthesis ~ Salt * N + Block + (1\|Fam) | 10 | 585.4 | -7.4 |
| Photosynthesis ~ Salt * N + (1\|Fam) | 6 | 580.4 | -12.4 |
| Photosynthesis ~ Salt * N | 4 | 578.4 | -14.5 |
| **Survival (N=1149)** |  |  |  |
| SNG ~ Salt + Pop + (1\|Fam) + (1\|Block) | 9 | 561.32 | 0 |
| SNG ~ Salt + Pop + (1\|Fam) | 8 | 559.32 | -2 |
| SNG ~ N + Pop + (1\|Fam) + (1\|Block) | 9 | 562.19 | 0.86 |
| SNG ~ N + Pop + (1\|Fam) | 8 | 560.19 | -1.14 |
| SNG ~ Pop + (1\|Fam) + (1\|Block) | 8 | 560.28 | -1.04 |
| SNG ~ Pop + (1\|Fam) | 7 | 558.28 | -3.04 |
| SNG ~ (1\|Fam) | 2 | 560.56 | -0.76 |

To gain insight about variance explained by models, we calculated the *R^2^* approximations proposed by (Nakagawa and Schielzeth, 2013). As these authors explain, in the context of GLMMs this leads to two different sorts of *R^2^*, a marginal *R^2^* that reflects variance explained by fixed factors only, and a conditional *R^2^* that reflects variance explained by both fixed and random factors. We report each of these as appropriate, e.g., where the best model includes only fixed factors we report only the marginal *R^2^*.

We assessed three survival states that were coded as 0 for live plants, 1 for dormant plants, and 2 for plants that died during the experiment. We modeled survival using random effects logistic regression. In one set of models, we included dormant plants as alive, and in another we excluded them. The results were qualitatively similar, so we report only the case where dormant plants were treated as alive.

## RESULTS

### Treatment validation

To ensure that our N amendment treatments were not flushed out during the once weekly flow through watering we measured the total dissolved N in leachate from a subsample of the seedlings. We found that N was not significantly different between the low salt-no N and the high salt-high N amended plants and, therefore, confirmed that we did not lose the N due to watering between treatments (Mean Square = 0.11, F ndf 3/ddf 48 = 0.23, Pr(>F) = 0.88).

### Trait responses to treatment

The only useful predictor among the fixed effects for the change in height is the height at the start of treatments in October (Table 1; marginal R^2^ = 0.82; Anova chisq= 4432.67; p <0.0001). The conditional *R*^2^ is 0.92, and the marginal *R*^2^ is 0.82, indicating that 82 % of the variance in change in height is explained by a positive relationship with the fixed effect of height at the beginning of the experimental treatments (estimate = 0.9009). The variance components (scaled to sum to 1) for population and maternal families within populations are 23% and 37% of the random variance respectively. A plot of the random effects suggest that the sites are largely similar, with the exception of FD which is very different (Figure S1). It’s not obvious that any population has more among-family variability, but our design is limited to determine this.

For LMA (log link), the best model included only the random effects of block, population and maternal families within populations. The conditional *R*^2^ is 0.99. In this model, maternal family explains 51% of the variance, population explains 30% of the variance and block explains 17% (Figure S2).

For succulence (log link) the best model included the fixed effect of salt and the random effects for family nested within population. The marginal *R^2^* = 0.39, while the conditional *R^2^* = 0.99. The scaled variance components for maternal families within populations and populations are, respectively, 0.91 and 0; in other words there is very large variance among maternal families within populations, but not among populations (Figure S3). While increased salt resulted in a statistically meaningful reduction in succulence (Figure 2), it is only when conditioned on family that it adds to the explanatory power of the model.

**Figure 2.**
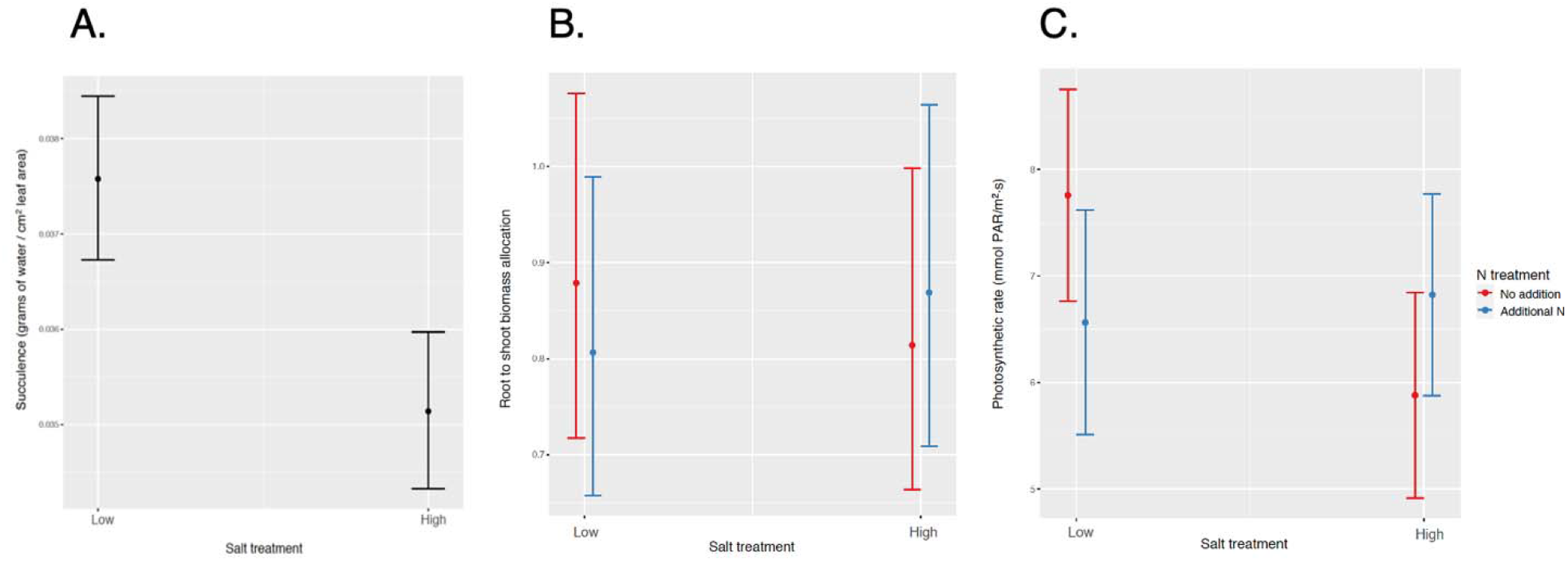
A) Succulence response to salt across populations with no N effect, including the random effects of maternal families within populations and populations (N=818). B) root to shoot biomass and C Photosynthetic rate response to N depends on salt treatment across all mangrove propagules. The best model for photosynthetic rate (N=118) does not include any of the random effects. The best model for root to shoot biomass (N=1073) also includes the random effects of maternal families within populations, populations and block.

The best model for total biomass included only the random effects of populations and maternal families within populations. The conditional *R^2^* = 0.15. The terms for populations and families nested within them account for 14% and 23% of the random variance, respectively (Figure S5).

Root to shoot biomass ratio was the only variable for which the data supported the saturated model as best model (Figure 2). The conditional *R^2^* = 0.134, while the marginal *R^2^* = 0.007. An Anova to evaluate these fixed effects revealed that the main effects of salt and N were not individually significant but the interaction was (chisq = 7.24; p= 0.007). However, the small size of the marginal *R^2^* suggests that these effects are mainly meaningful when conditioned on the random terms. The random terms population, population and block account for 12%, 18%, and 10% of the random variance, respectively. For this trait, population WI was different from all the others (Figure S4).

The photosynthesis data were most limited in sample size since we were only able to assess one individual with at least two healthy leaves for each surviving maternal line for each treatment (N=118). The best model was one including the fixed effects salt, N, and their interaction, but no random effects (Table 1). As with root to shoot biomass ratio, an Anova to evaluate these fixed effects showed the main effects of salt and N were not significant but the interaction was (chisq = 4.46; p = 0.035). At ambient N levels (no N added), maximum photosynthetic rate and root-to-shoot ratio both declined with increasing salt concentration, but this negative impact of salt was absent or reversed upon the addition of N fertilizer (Figure 2B and 2C).

In summary, we found that some combination of salt and N treatments were included in the best models for three of the six traits we measured. Succulence decreased in response to higher salt (Figure 2A), but this response varied largely by family (Figure S3). For root to shoot biomass ratio and maximum photosynthetic rate the responses to experimental treatments depended on changes in both salt and N (Figure 2B and 2C). Root to shoot ratio also varied by family and population (Figure S4).

### Survival

Of 1149 plants, 76 were unequivocally dead in April, 1073 unequivocally alive, and 166 dormant. The number of plants that showed active growth ranged from 63% in HI to 91% in WI (Figure 3). The number of plants that didn’t show growth, but also didn’t appear to be dead ranged from 3% in WB and WI to 7% in UTB. We modeled survival by including these dormant plants alternatively as either alive or dead. In both cases the best-supported model was one including a fixed effect for population and a random effect for maternal family (Figure S7). The model treating dormant plants as alive had conditional R^2^ = 0.11 and marginal R^2^ = 0.08. The model with dormant plants as dead had conditional R^2^ = 0.28 and marginal R^2^ = 0.13. Because there is no residual in the equation defining logistic regression, no variance component for a random residual is calculated, and thus it is not possible to calculate a meaningful scaled variance component for family here.

**Figure 3:**
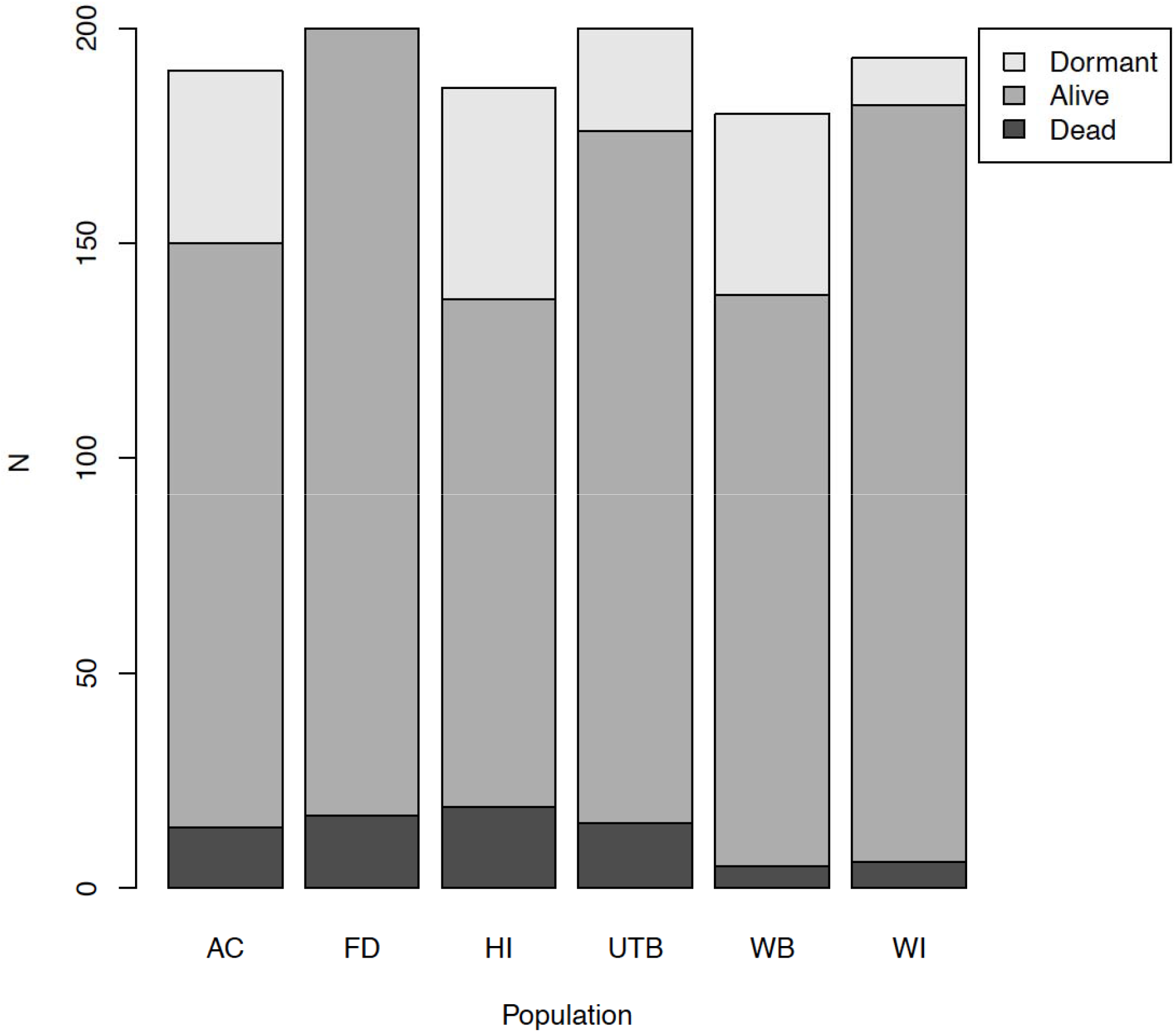
Absolute numbers of plants in each growth state at harvest by population of origin. Dormant plants (light gray) showed no signs of desiccation or growth. Seedlings from the HI population had the lowest survival overall, while WI seedlings had the highest survival. Plants alive at harvest are medium gray, plants that had died by harvest are dark gray.

## DISCUSSION

We assessed the growth and survival of *R. mangle* propagules to full factorial combinations of two salt and two N levels because these are two important abiotic properties that potentially have important impacts on mangrove biomass and traits and, by extension, on the biodiversity and ecosystem function of coastal wetlands. In addition to natural variation in these conditions, anthropogenic activities may result in more extreme levels of these conditions from runoff and flooding (Antlfinger and Dunn, 1983; Ellison et al., 2005; Krauss et al., 2006; Gedan et al., 2009; Kirwan and Megonigal, 2013; Lewis et al., 2014). We predicted that *R. mangle* seedlings would be plastic in response to salinity and N amendment in putative adaptive traits that conserve water and adjust allocation of N. Our study showed that only succulence was plastic in response to salt, regardless of N treatment. Allocation to root and shoot biomass and maximum photosynthetic rate were also plastic, but response of both traits to N amendment depended on the level of salt. This supported our second prediction that response to salinity and N amendment will co-vary as plants shift resources to maintain osmotic balance. We also found support for our third prediction that populations would vary in putative adaptive traits, and in plasticity of these traits, due to population differentiation. Importantly, every trait except for photosynthesis varied among population and maternal families within populations. This was also true of survival. In fact, maternal family and population were the most consistent predictors for variation in traits and survival.

### Phenotypic plasticity in response to treatments

We expected that *R. mangle* propagule traits and survival would respond to salinity by decreasing growth and respond to N fertilization by increasing growth. We also expected that N could alleviate some of the effects of salinity as indicated by an interaction of the two conditions. However, we found no response to treatments in height growth, succulence or total dry biomass. This may be due to the propagules being supported by resources provided by the maternal tree, which in *R. mangle* can support growth for at least a year (Ball, 2002; Proffitt and Travis, 2010). If the seedlings were supported by these maternal reserves, height growth would likely be more correlated to propagule length at collection which would be corrected for in the start-of-experiment (time zero) height measurements that we included as a covariate. Because our treatment duration was only six months, the lack of growth response to treatments is consistent with dependence on maternal reserves. However, we did see seedling response to treatment in succulence, maximum photosynthetic rate and allocation to root and shoot biomass.

Increased succulence is a common response to water deficiency under high salinity conditions (Rosenthal et al., 2002; Vendramini et al., 2002; Ottow et al., 2005; Karrenberg et al., 2006; Richards et al., 2008, 2010), but in our experiment we detected reduced succulence in response to high salt. However, this could be due to the fact that *R. mangle* excludes salt, which may result in a different physiological response to salinity (Cavalieri and Huang, 1979; Donovan et al., 1996; Tester and Davenport, 2003). For example, several other halophytes that are salt excluders, including the succulent plant *Salicornia europea* L., and another member of Rhizophoraceae, *Kandelia candel* (L.) Druce, do not increase succulence or leaf thickness in response to high salinity (Glenn and O’Leary, 1984; Kao et al., 2001). In fact, with N fertilization *K. candel* decreased leaf thickness when salinity was increased (Kao et al., 2001). Thus, one possible explanation for our results is that the N fertilized seedlings were able to reallocate resources and still maintain turgor and water uptake in the high salt condition without increased succulence.

Although we saw significant plasticity for three of the six traits in response to our treatments, response in only R:S and maximum photosynthetic rate depended on the interaction between salt and N fertilization. We expected maximum photosynthetic rate to increase in response to N fertilization because the enzyme RuBisCO, which catalyzes the dark reactions in photosynthesis, requires a large amount of N (Sage et al., 1987; Andersson, 2008). Further, a meta-analysis across different species and biomes showed increased maximum photosynthetic rate with increased N (Walker et al., 2014). This could lead to increased shoot biomass which is supported by our analysis in the absence of salt (i.e. decreased R:S in response to N). Despite this expectation, there was no overall response to high N level independent of salt treatment. One reason might be that photosynthesis was limited by other nutrients, not just N, and thus increasing N alone might not have been enough to elicit a response. In a field study, dwarf *R. mangle* did not respond to N alone, but did increase biomass in response to fertilization with N, phosphorus (P), and potassium ions (K^+^), potentially because they were P limited (Feller, 1995). We also expected that response to N amendment would depend on salinity. We found that in plants treated with high salt, maximum photosynthetic rate was slightly enhanced by high N. Possibly, the additional N enabled the plants to synthesize N-rich compatible solutes for osmotic regulation and continue photosynthetic gain of carbon (Flowers and Colmer, 2008). Plants in high salt also responded differently in allocation with increased N; instead of increasing shoot biomass, they increased root biomass on average.

### Variation within and among sites

Phenotypic variation that is maintained in common garden from within and among populations would indicate *R. mangle* has heritable trait diversity to adapt to changing environmental conditions. We found variation in height growth, succulence, R:S, LMA, and total dry biomass was largely determined by maternal families within populations. Seedling survival depended on population and varied among maternal families for all of the six populations. Proffitt and Travis (Proffitt and Travis, 2010) also found seedling survival varied among maternal families, as well as by location in the intertidal zone. But in their study after three years, growth and survival did not reflect initial propagule size (Proffitt and Travis, 2010). Our results support these previous findings that propagule length is positively correlated to short term performance, which suggests that maternal reserves in the *R. mangle* propagule can help the seedling survive, and larger propagules contain more maternal reserves than smaller propagules (Ball, 2002; Proffitt and Travis, 2010). Because our study was a short-term, controlled greenhouse study, maximum photosynthetic rate might be the best indicator for an immediate response. Variation in maximum photosynthetic rate can ultimately manifest as variation in growth and allocation of resources, particularly once the seedling has depleted maternal reserves. The seedlings did not show significant differences in height growth or total dry biomass in response to treatments. But given the plasticity we saw in maximum photosynthetic rate, and the one-year and three-year growth results found in a previous study of nearby populations (Proffitt and Travis, 2010), it’s conceivable that with additional time our seedlings would respond more dramatically to these treatments. However, previous work in several different systems have argued that greenhouse studies are often unable to recreate relevant field conditions so these responses may only be registered in field conditions (Schittko et al., 2016; Rinella and Reinhart, 2017; Forero et al., 2019; Dostál et al., 2020).

The amount of heritable phenotypic diversity and differentiation we discovered in this study is an important indicator of the potential for this species to respond to changing conditions, which may be surprising. We previously reported low genetic diversity among these plants based on molecular markers, which is expected to limit the potential for different responses among individuals. On the other hand, we discovered high epigenetic diversity (Mounger et al., 2021), which could contribute to phenotypic and functional diversity, and could be a mechanism underlying the type of phenotypic differences and plasticity we found here (Zhang et al., 2013; Nicotra et al., 2015; Herrera et al., 2017; Jueterbock et al., 2020). In addition, we know very little about the interactions with the microbiome in the species, but microbes have been highlighted as important symbionts in these and other challenging environments (Bowen et al., 2017; Angermeyer et al., 2018; Jung et al., 2021). Soil microbial activity could have dramatic impacts on the future nutrient availability and stability of these coastal sediments (Deegan et al., 2012; Bowen et al., 2017; Hughes et al., 2020; Lewis et al., 2021). In fact, a recent study suggested that bacterial community composition differed among *R. mangle* maternal genotypes but not with genetic diversity (Craig et al., 2020).

## Conclusions

Mangroves provide many ecosystem services, but their global area has declined between 20% and 35% since 1980 (Food and Agriculture Organization of the United Nations, 2007; Global forest resources assessment 2020, 2020). In addition to global area decline due to anthropogenic activities, mangroves also face rising sea levels and flooding predicted by climate change (Krauss et al., 2008; IPCC, 2014; Wuebbles et al., 2014; Osland et al., 2017). Previous work suggested that *R. mangle* growth and survival depended on an interaction of intertidal elevation and maternal genotype interaction, suggesting variation in response to flooding conditions (Proffitt and Travis, 2010). This variation among genotypes could be important to enable *R. mangle* to dominate over a larger intertidal range. However, in addition to changes in flooding, anthropogenic activities are causing changes in salinity and N level in *R. mangle* ecosystems (McKee et al., 2007). The interacting effects of salinity, N level, and elevation are complex and potentially non-additive (McKee et al., 2007). Our experimental findings suggest that the important traits of succulence, photosynthesis and root to shoot biomass allocation respond to salinity, N level, or the combination of these conditions, but also the magnitude of responses varies among populations and even maternal families within populations. In addition, *R. mangle* seedling survival depended on maternal family for all of the six sites.

This variation in important traits and survival among families and among populations is particularly interesting given our previous work with genetic markers that showed that these populations had low genetic diversity, and little population differentiation. Considering the importance of this foundation species for the functioning of the coastal ecosystem, the lack of genetic diversity might be alarming. However, accumulating studies provide important evidence that genetic variation must be interpreted with caution (Hufford and Mazer, 2003) and that the emphasis on only variation in DNA sequence can be misguided (Keller, 2002, 2014; Sultan, 2015; Bonduriansky and Day, 2018). Nongenetic sources of response may contribute to the resilience of *R. mangle* and other critical species to changing environmental conditions and contribute to future adaptation to a complex mosaic of environmental conditions.

## DATA AVAILABILITY STATEMENT

All data and scripts will be submitted on zenodo and updated here as soon as possible.

## AUTHOR CONTRIBUTIONS

CLR, KLL, GAF, and DBL conceived the study, and designed the experiments. KLL and JM collected plants and maintained experiments. KLL and GAF analyzed the data. CLR and KLL wrote the first draft of the manuscript. All co-authors provided input and revisions to the manuscript.

## ACKNOWLEDGMENTS

We thank Bert Anderson, Sandy Voors, Viviana Penuela Useche, M. Teresa Boquete, Mariano Alverez, and Marta Robertson for their help with analyses and review of earlier versions of this manuscript. We are grateful to Christine Brubaker, Racquel Pancho, Brianna Jerman, Vernetta Williams, Alan Franck, and Mary Mangiapia for their guidance, and Charley’s Pizza for great food during greenhouse and plant processing work. We thank Samantha Blonder, Jordan Dollbaum, Maria Nikolopoulos, Shane Palmer, Bradley Biega, Bryan Lotici, Harper Cassidy, Jelena Dosen, Nancy Sheridan, and Dawei Tang for their help in the field and the greenhouse. We thank our friends and family for constant support and encouragement and we acknowledge support by the Open Access Publishing Fund of University of Tübingen.

## Supplementary Material

### Supplemental figures

**Figure S1.**
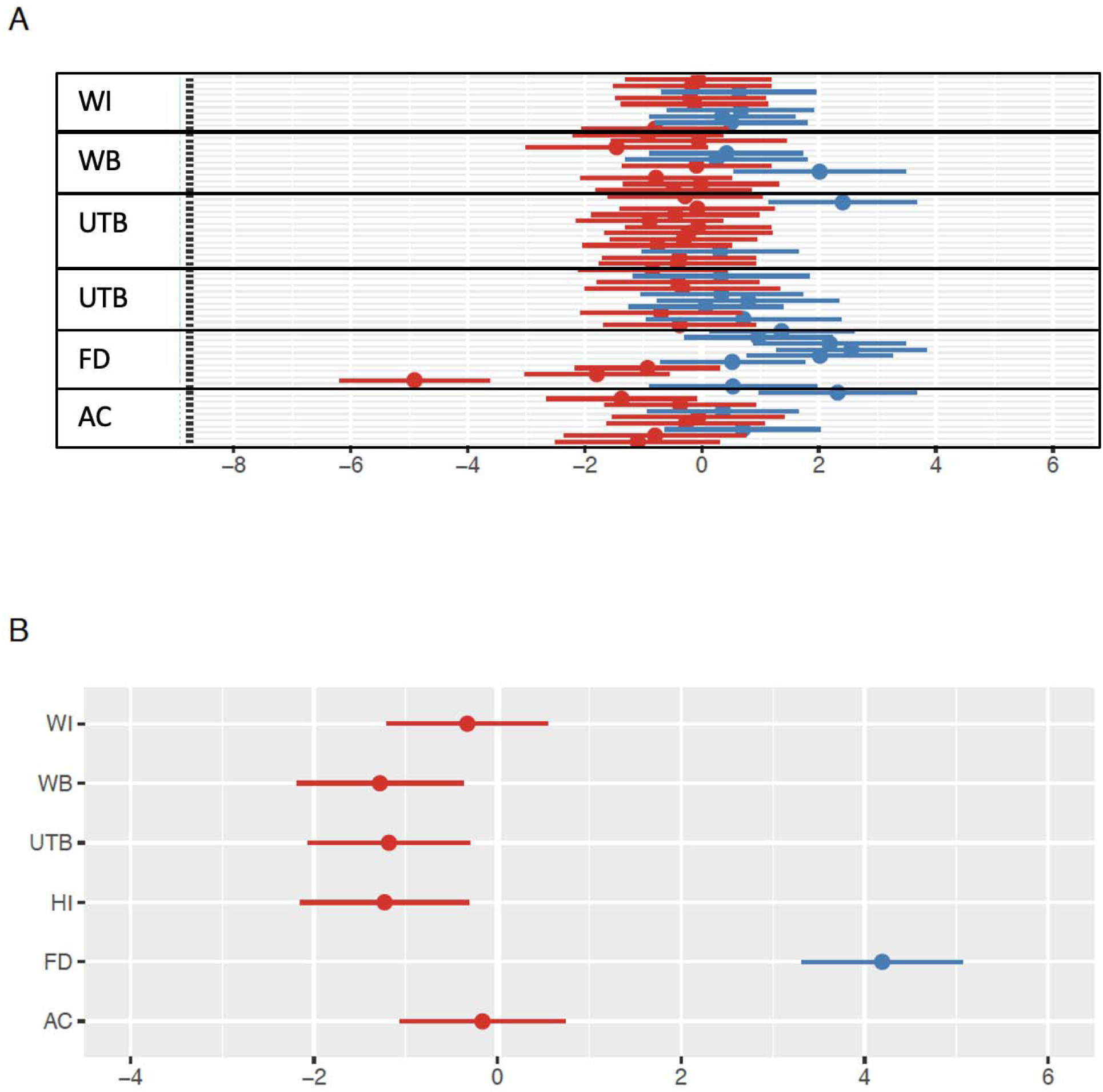
Random effects of (A) maternal families and (B) populations on total height.

**Figure S2.**
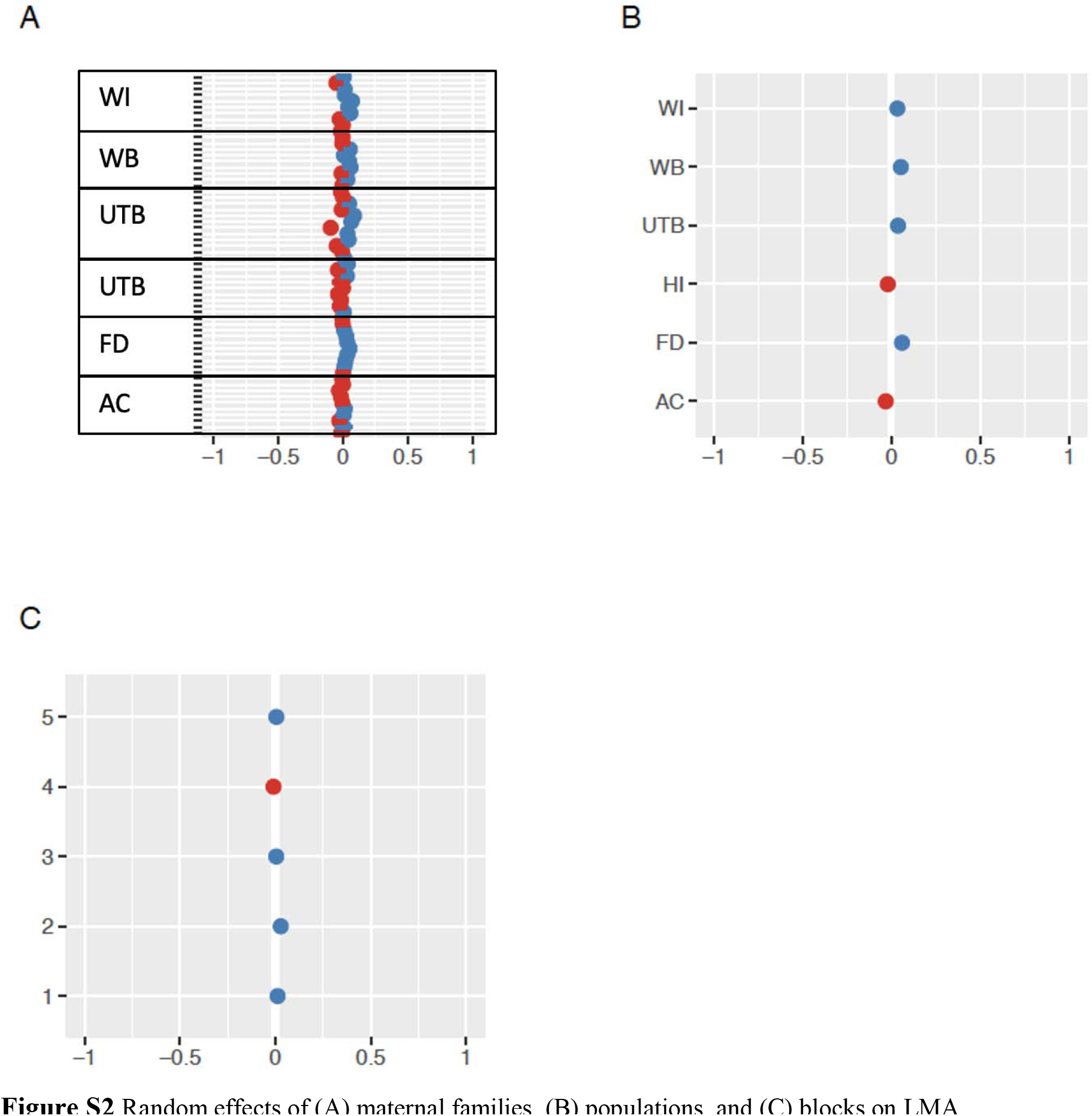
Random effects of (A) maternal families, (B) populations, and (C) blocks on LMA.

**Figure S3.**
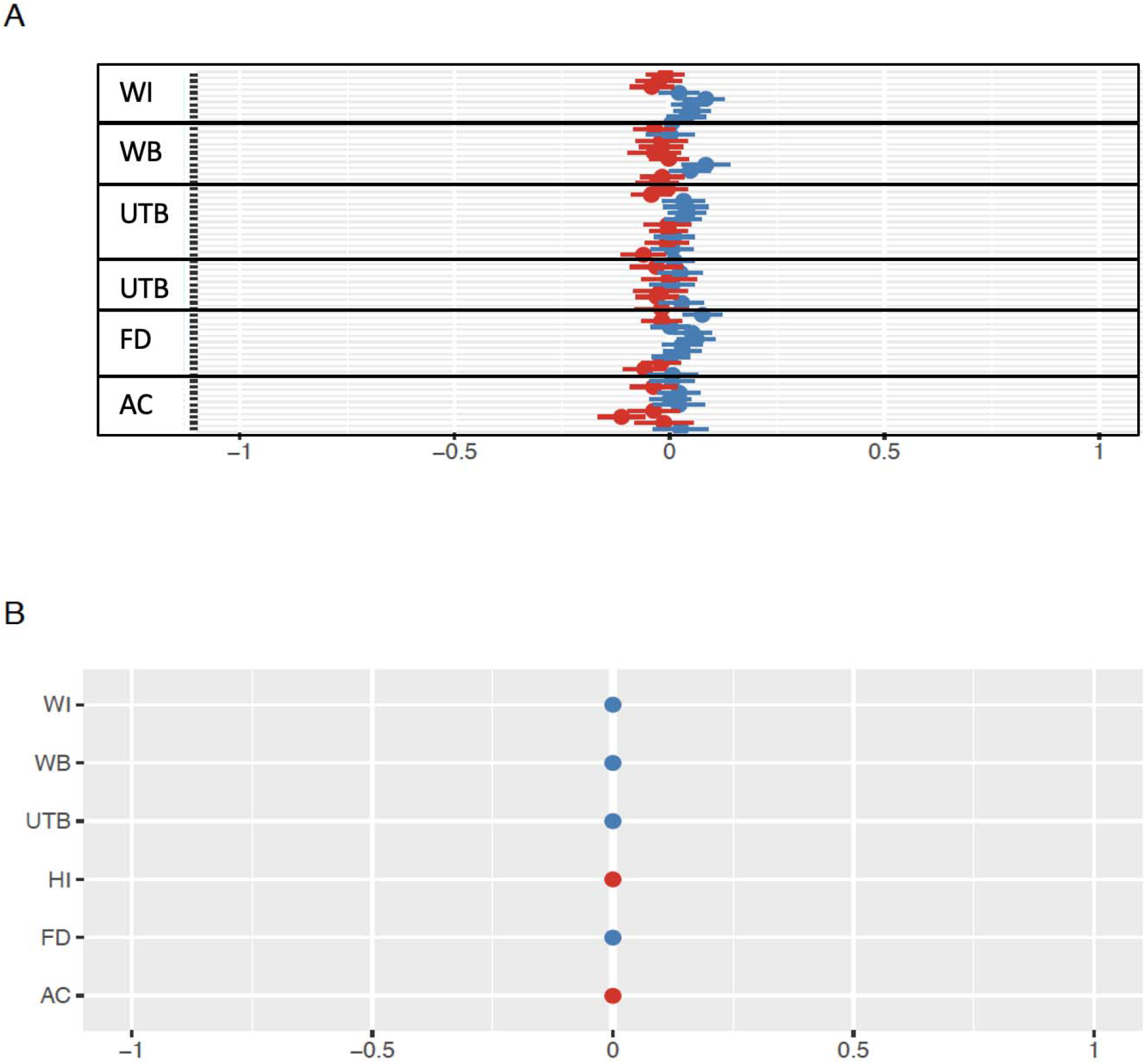
Random effects of (A) maternal families and (B) populations on total succulence.

**Figure S4.**
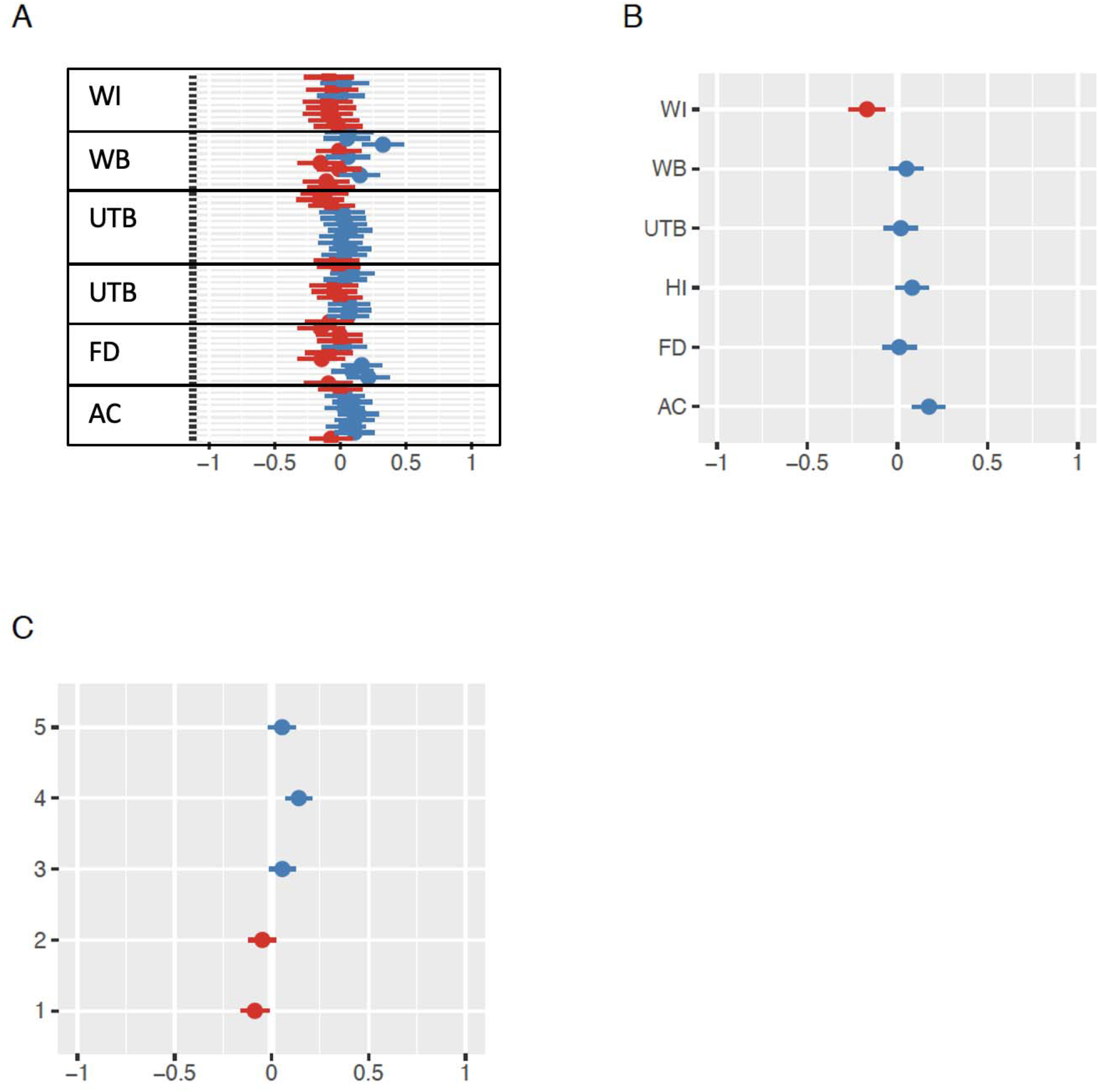
Random effects for (A) maternal families, (B) populations, and (C) blocks, in the root-shoot model.

**Figure S5.**
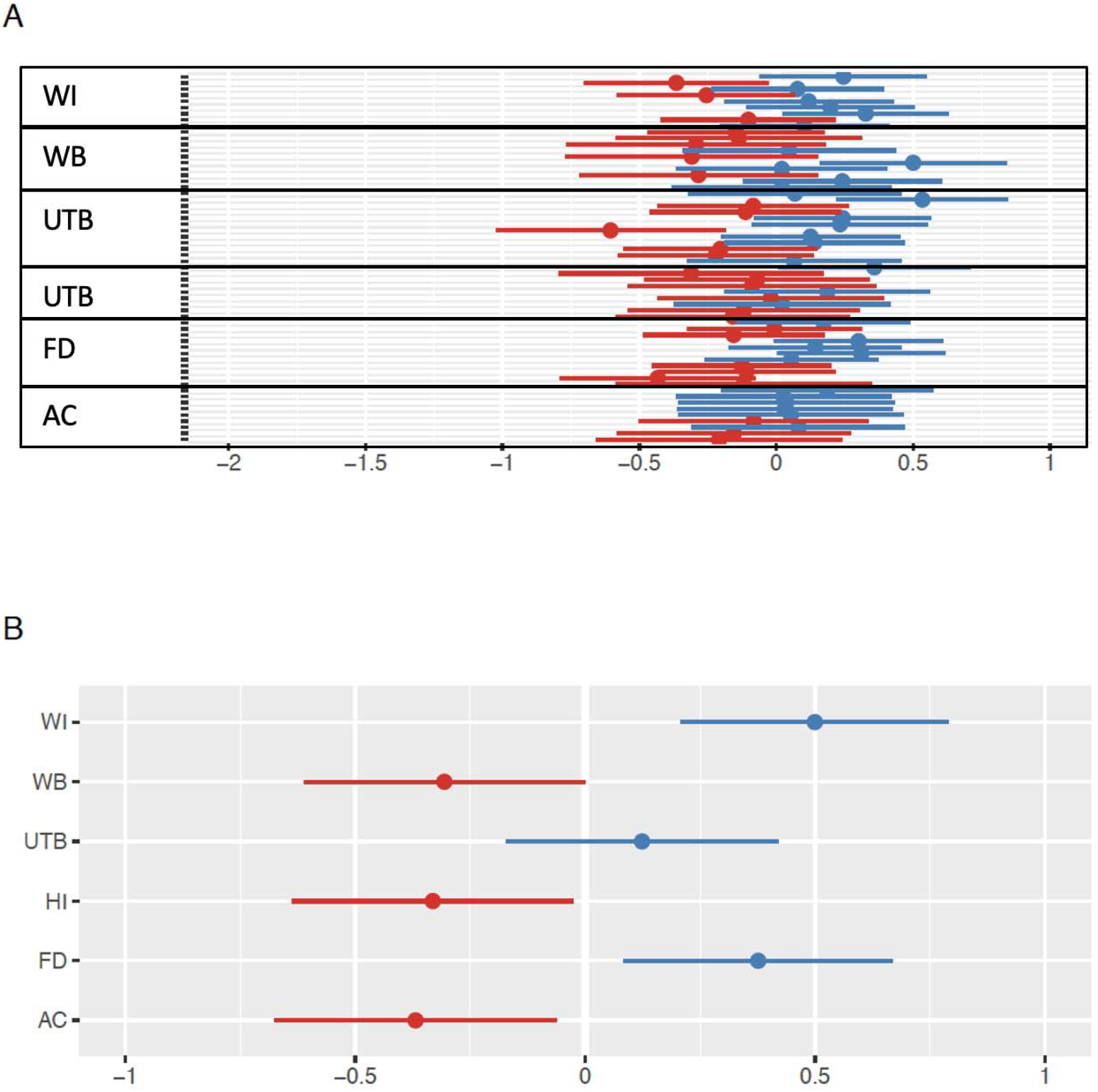
Random effects of (A) maternal families and (B) populations on total biomass.

**Figure S6.**
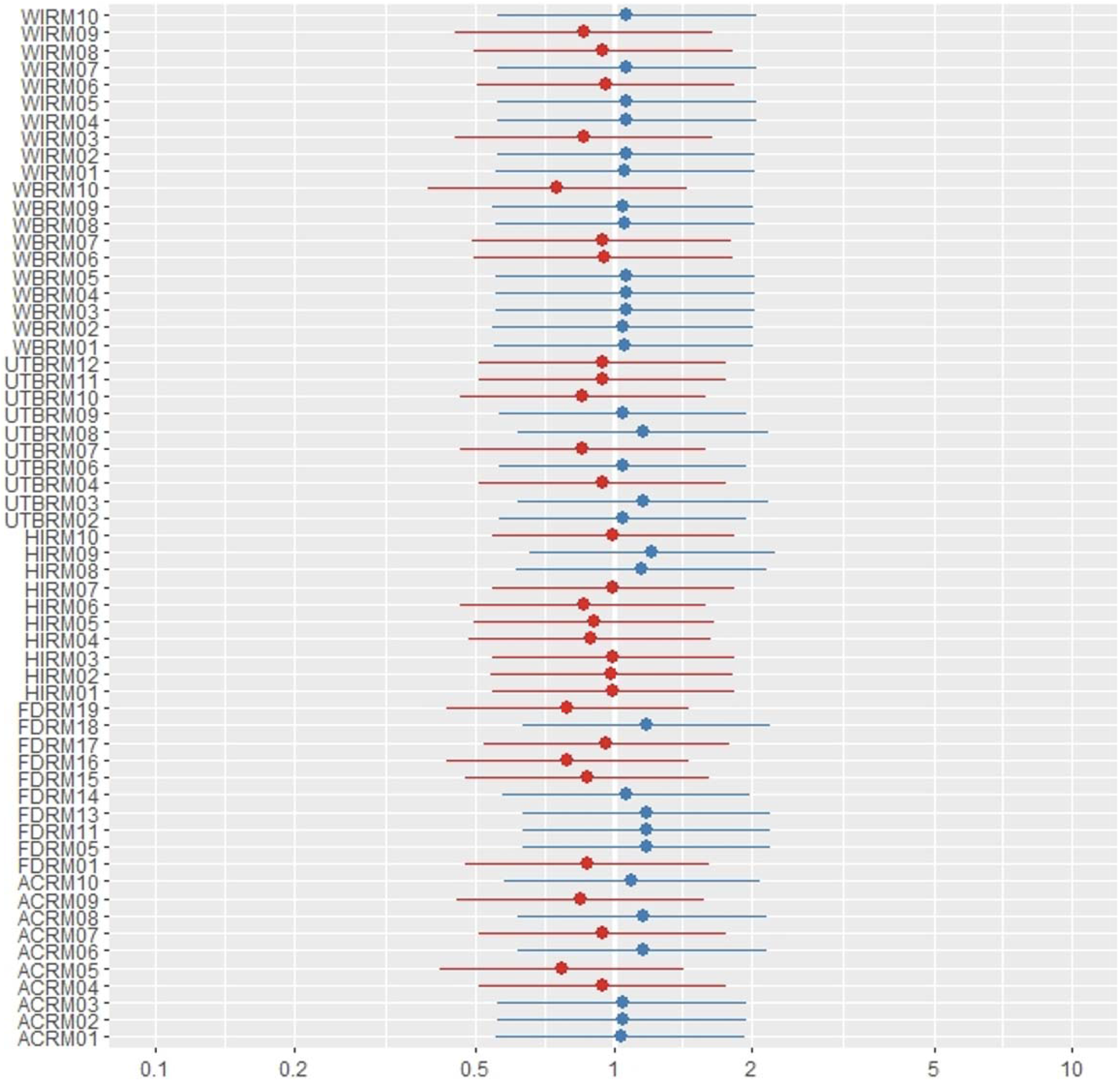
Random effects of maternal families on survival.

